# The pathogenic p.N1662D SCN2A mutation reveals an essential molecular interaction for Na_v_1.2 channel inactivation

**DOI:** 10.1101/2023.11.12.566785

**Authors:** Géza Berecki, Elaine Tao, Katherine B. Howell, Rohini K. Coorg, Kris Kahlig, Markus Wolff, Ben Corry, Steven Petrou

## Abstract

Mutations in the SCN2A gene encoding the Na_v_1.2 sodium channel can lead to neurodevelopmental disorders. We studied the N1662D variant associated with severe early-onset developmental and epileptic encephalopathy (DEE). The N1662D mutation almost completely prevented fast inactivation without affecting activation. The comparison of wild-type and N1662D channel structures suggested that the ambifunctional hydrogen bond formation between residues N1662 and Q1494 is essential for fast inactivation. Fast inactivation could also be prevented with engineered Q1494A or Q1494L Na_v_1.2 channel variants, whereas Q1494E or Q1494K variants resulted in incomplete inactivation and persistent current. Molecular dynamics simulations revealed a reduced affinity of the hydrophobic IFM-motif to its receptor site with N1662D and Q1494L variants relative to wild-type. These results demonstrate that the interactions between N1662 and Q1494 underpin the stability and the orientation of the inactivation gate and are essential for the development of fast inactivation. Six DEE-associated Na_v_1.2 variants, with mutations mapped to channel segments known to be implicated in fast inactivation were also evaluated. Remarkably, the L1657P variant also prevented fast inactivation and produced biophysical characteristics similar to N1662D, whereas the M1501V, M1501T, F1651C, P1658S, and A1659V variants resulted in biophysical properties that were consistent with gain-of-function and enhanced action potential firing of hybrid neurons in dynamic action potential clamp experiments. Paradoxically, low densities of N1662D or L1657P currents potentiated action potential firing, whereas increased densities resulted in sustained depolarization. The contribution of non-inactivating Na_v_1.2 channels to neuronal excitability may constitute a novel cellular mechanism in the pathogenesis of *SCN2A*-related DEE.

**SIGNIFICANCE STATEMENT:** *SCN2A* gene-related early-onset developmental and epileptic encephalopathy (EO-DEE) is a rare and severe disorder that manifests in early infancy and childhood. *SCN2A* mutations affecting the fast inactivation gating mechanism can cause altered voltage dependence and incomplete inactivation of the encoded Na_v_1.2 channel, leading to abnormal neuronal excitability. In this biophysical and clinical study of neuronal Na_v_1.2 variants, we identified amino acid residues that are critical for the stability and orientation of the inactivation gate during fast inactivation. Mutations of these residues prevent fast inactivation and may lead to EO-DEE via a novel pathophysiological mechanism. The results provide novel structural insights into the molecular mechanism of Na_v_1.2 channel fast inactivation and inform treatment strategies for *SCN2A*-related EO-DEE.

## INTRODUCTION

The *SCN2A* gene-encoded voltage-gated Na_v_1.2 channels are predominantly expressed in the excitatory neurons of the central nervous system, where they contribute to the generation and propagation of action potentials. Na_v_1.2 channels activate in response to depolarizing stimuli to allow the influx of Na^+^ ions. Activation is followed by fast inactivation, a mechanism which terminates the Na^+^ conductance even in the continued presence of depolarization. This mechanism is important for controlling neuronal excitability by preventing rapid refiring and allowing the membrane potential to return to resting values once the signal has been sent.

Fast inactivation of sodium channels has been intensely studied and reviewed for nearly 80 years (1–7). Channel activation is initiated by the rapid movement of segment 4 (S4) voltage sensors in domains I-III (S4_DI-III_) leading to pore opening followed by the slower movement of S4_DIV_ (8), which represents the rate-limiting step for the development of fast inactivation and the recovery from this state (9). A sequence of three conserved hydrophobic amino acids (IFM motif) in the DIII-IV linker (inactivation gate) is essential for fast inactivation since mutations of these residues are able to slow or completely remove fast inactivation (10). In rat brain sodium channels, alanine scanning mutagenesis of the S4-5_DIV_ region and subsequent two-microelectrode voltage clamp experiments showed incomplete fast inactivation with F1651A or L1660A channels, and a nearly abolished fast inactivation with the N1662A mutant relative to wild-type (11). It has been suggested that the mutated residues form part of the inactivation gate’s IFM motif receptor (3, 11).

The availability of sodium channel structures provides a unique opportunity to further unravel the fast inactivation mechanism at molecular level (12–16). This process involves a series of electrostatic interactions between amino acid residues located on S4_DIV_, S4-5_DIV_ linker, DIII-IV linker (inactivation gate), S6_DIII_ and S6_DIV_ of the intracellular pore module, and the C-terminal domain (CTD) (12). A hydrophobic receptor site mapped to residues in the S4-5_DIII_ and S4-5_DIV_ linkers and outside of the helices lining the activation gate represents the IFM motif binding site (12, 15, 17). Recent data suggest that in addition to IFM motif binding, the closure of two hydrophobic rings at the bottom of S6 helices is also needed to complete fast inactivation (18). However, despite the experimental data and the novel cryo-EM structures of various neuronal Na_v_ channels (13, 14, 19, 20), fundamental questions regarding the precise molecular interactions mediating Na_v_1.2 fast inactivation still prevail.

De novo missense mutations in *SCN2A* can lead to neurodevelopmental disorders of various severity, including early-onset developmental and epileptic encephalopathy (DEE), a rare severe disorder caused by Na_v_1.2 gain-of-function (21–23). Early-onset DEE-related Nav1.2 channel variants often show impaired biophysical properties due to a changed voltage dependence, time course and/or extent of fast inactivation. Patients are at high risk of premature mortality from direct and indirect effects of seizures, which typically beginning within days of birth, and are often difficult to control with antiseizure medications. Understanding the mechanisms of ion channel dysfunction leading to abnormal neuronal excitability should facilitate the interpretation of the neurological disease and the implementation of mechanism-targeted therapies.

In this study, we assessed the biophysical and the structural consequences of the N1662D missense mutation identified in a patient with severe early-onset DEE. This mutation is predicted to cause local structural changes that severely alter fast inactivation and neuronal excitability. We used engineered Na_v_1.2 channel variants to probe the role of hydrogen bond formation between residues N1662 (located in the cytoplasmic end of S5_DIV_), and Q1494 (located in the DIII-IV linker) in fast inactivation. Similarly, we also assessed six DEE-associated Na_v_1.2 missense mutations located nearby the N1662 residue (L1657P, P1658S, A1659V), the S4-5_DIV_ linker (F1651C) or the DIII-IV linker (M1501V, M1501T) to determine their impact on fast inactivation and neuronal excitability. To better understand the structural changes caused by the N1662D mutation relative to wild-type, we used molecular dynamics (MD) simulations and evaluated the binding stability of the IFM motif and other key interactions between residues in the S4-S5_DIV_ linker and DIII-IV linker. The results suggest that the N1662-Q1494 interaction plays a critical role in maintaining a stably bound inactivation gate and mutations of either residue lead to disrupted fast inactivation.

## RESULTS

### The N1662D mutation disrupts fast inactivation

The N1662D mutation has been identified by whole exome sequencing in a patient who presented with severe early-onset DEE (24). A summary of the phenotypic data for the patient with the N1662D variant is shown in SI Appendix.

The N1662 residue is located at the cytoplasmic end of S5_DIV_, near the inactivation gate (DIII-IV linker) and the S4-5_DIII_ and S4-5_DIV_ linkers, in the proximity of the intracellular gate (Figure 1) (13). This suggest that the N1662D mutation may severely affect fast inactivation, in agreement with previous alanine scanning mutagenesis data (11). The investigation of the wild-type channel structure revealed that the N1662 residue is involved in intermolecular interactions with several adjacent residues, including Q1494 and F1489 in the D_III-IV_ linker, and P1657 in the S4-5_DIV_ linker. Remarkably, the hydrogen bonds between the N1662 and Q1494 residues are simultaneous donor and acceptor interactions that may impact the stability and orientation of the α-helices (25). The N1662D mutation likely disrupts hydrogen bond formation between N1662 and Q1494 (Figure 1C).

**Figure 1.**
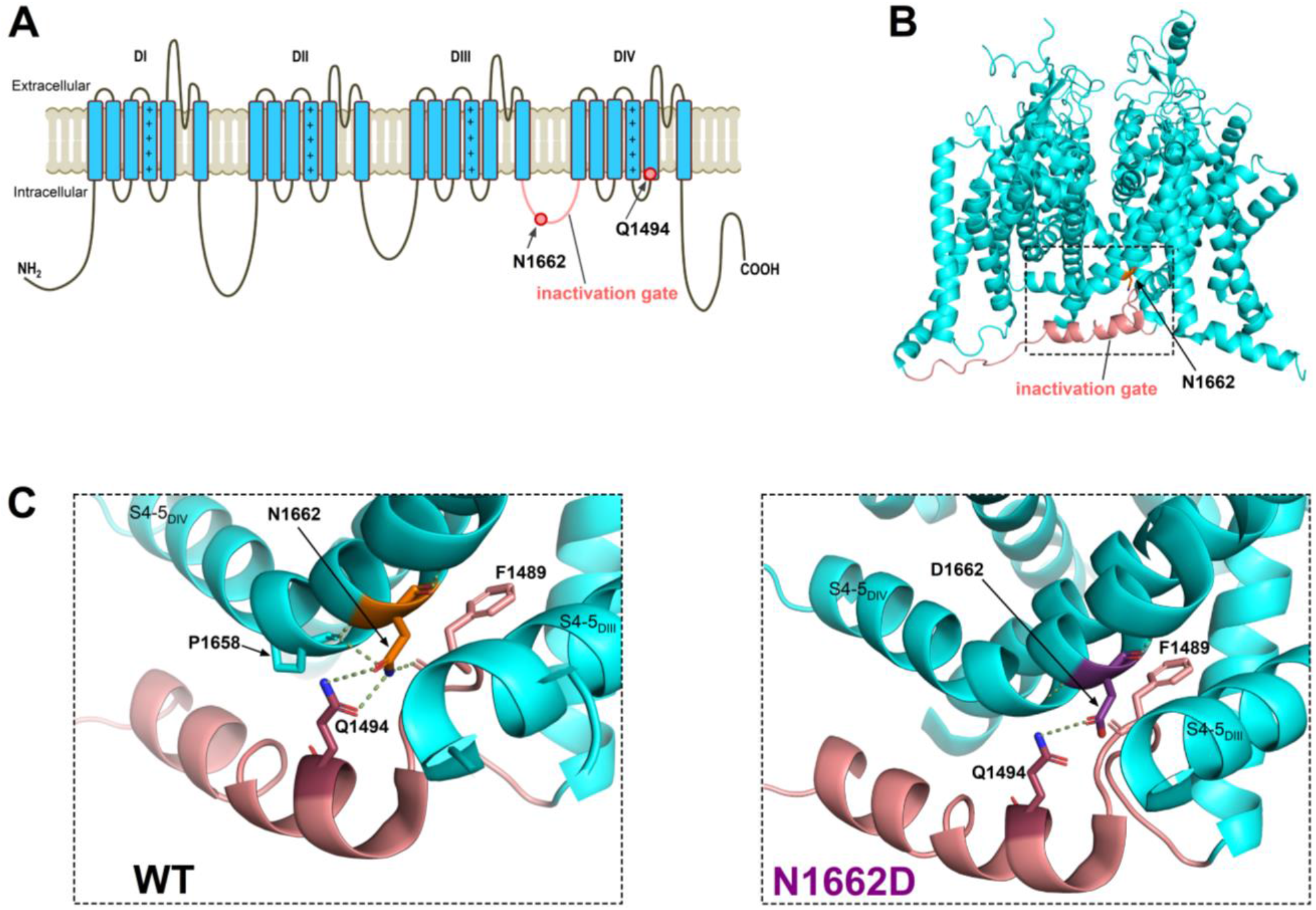
Location of amino acid residue 1662 and impact of N1662D mutation on polar interactions. **(A)** 2D **t**ransmembrane topology of the wild-type (WT) Na_v_1.2 channel showing domains DI−DIV and amino acid residues N1662 (cytosolic side of S5_DIV_) and Q1494 (DIII-DIV linker/inactivation gate, in pink). The positive charges in the S4 voltage-sensing segments are marked. **(B)** 3D side-view of the WT Na_v_1.2 channel (PDB accession no. 6J8E)(13). The boxed area highlights the N1662 residue and the inactivation gate. **(C)** Hydrogen bonds (dotted lines) between N1662 and Q1494 are ambifunctional (simultaneous donor and acceptor) in the WT channel (left). The N1662D mutations abolishes the hydrogen donor function of residue 1662 (right). Note that in the WT channel, N1662D also forms polar interactions with P1658 (located in S4-5_DIV_ and F1489 (IFM motif). The N1662 to Q1494 distance measured between the electronegative nitrogen amines (in blue) and carbonyl oxygens (in red) is 2.7 Å and 3.4 Å, whereas the D1662 to Q1494 distance is 3.4 Å.

To determine the impact of the N1662D mutation on the biophysical properties of the channel, we transiently expressed the wild-type or the N1662D Na_v_1.2 channel variants in Chinese hamster ovary (CHO) cells and recorded whole-cell sodium currents (I_Na_) using the voltage clamp technique. Relative to wild-type, the N1662D mutation caused a 5-fold reduction of the I_Na_ density and almost completely prevented inactivation, resulting in a relatively small decline of the inward peak I_Na_ amplitude at the end of 100-ms depolarizations (Figure 2A, 2B, Table 1). Relative to wild-type, the voltage dependence of N1662D activation was unchanged, however the voltage dependence of the inactivated N1662D current fraction exhibited a 12-mV shift towards depolarized potentials (Figure 2B, Table 1). Such positive shift of the inactivation curve may predispose to enhanced neuronal excitability (26). On the other hand, the lack of sodium channel inactivation could lead to a sustained depolarization of the membrane potential and prevent action potential firing in neurons (27, 28). Recovery from inactivation was faster for the N1662D variant relative to wild-type (Figure 2B, Table 1). This may reflect a reduced affinity of the inactivation gate receptor for the IFM motif due to a non-absorbing inactivation state from which recovery is faster (29).

**Figure 2.**
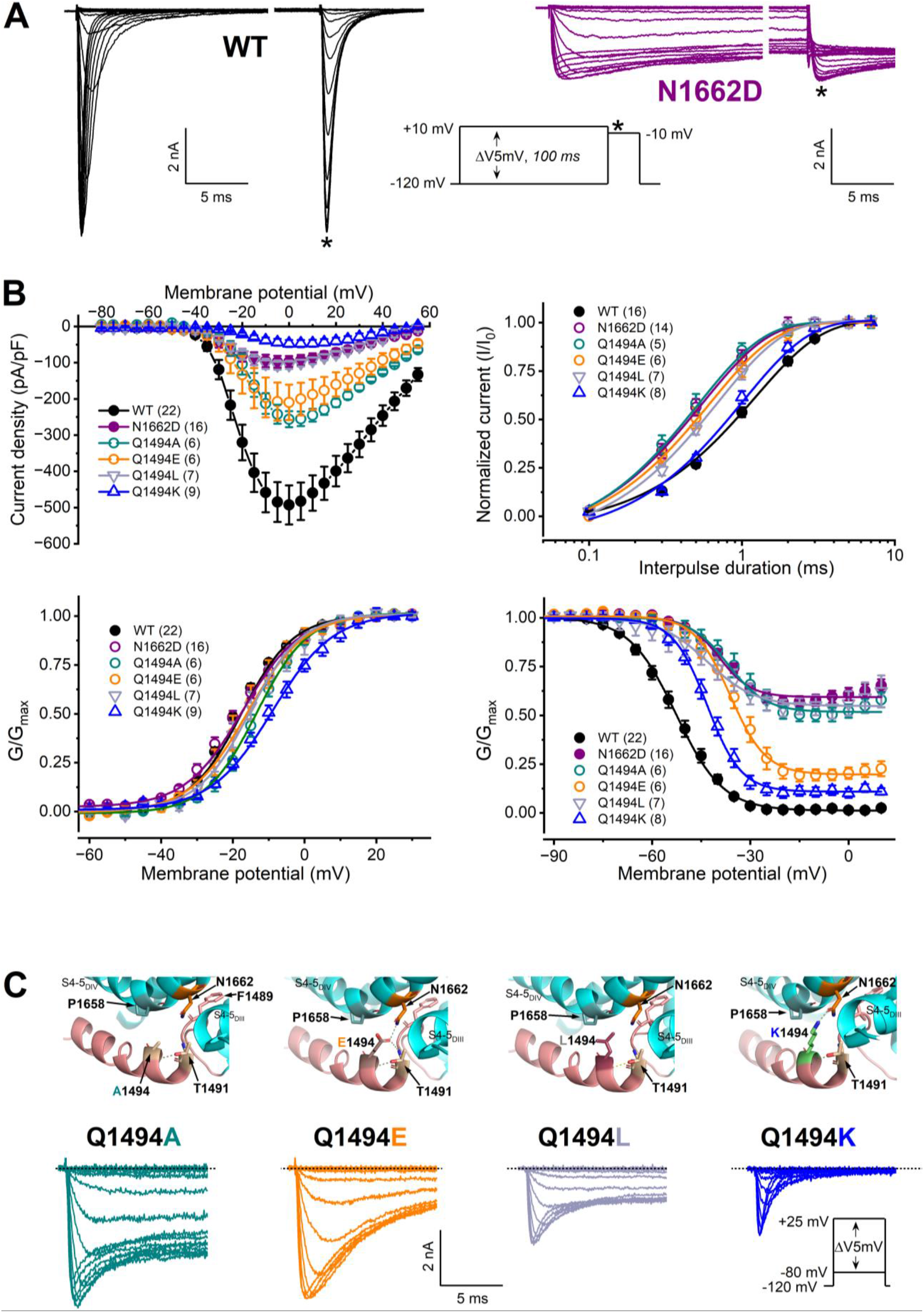
The N1662D mutation and inactivation gate residue Q1494 mutations prevent fast inactivation. **(A)** Representative wild-type (WT) and N1662D current traces elicited by 100-ms depolarizing (conditioning) voltage steps in 5 mV increment from a membrane potential of −120 mV, followed by a 20-ms test pulse to −10 mV to determine current availability (asterisk); inset protocol. For a better resolution, an approx. 80-ms break was implemented on the x-axis of the WT and N1662D traces, 15 ms after the onset of the inward current. **(B)** Current density-voltage relationships (top left). Time course of recovery from fast inactivation (top right). The time constants (τ values) of recovery were obtained by nonlinear fits of data to a single exponential equation (equation 2). Voltage dependence of activation (bottom left) and inactivation (bottom right). Activation was assessed using the inset protocol shown in C, whereas the protocol assessing steady-state inactivation is shown in A. The normalized conductance-voltage relationships are plotted as G/G_max_ values versus voltage and are referred to as activation or inactivation curves. Curves were obtained by non-linear least-squares fits of data to Boltzmann equations (equation 1, Materials and Methods). In all panels, the number of experiments, n, is shown in parentheses. The parameters of the fits and the statistical evaluation are included in Table 1. **(C)** Mutations of the inactivation gate residue Q1494 prevent or slow fast inactivation. Top: zoomed-in views of the S5_DIV_ and DIII-DIV linker (inactivation gate) residues, showing altered polar interactions between N1662 and A1494, E1494, L1494, or K1494 relative to Q1494 (WT, see Figure 1). Bottom: representative WT and mutant I_Na_ traces elicited from a holding potential of −120 mV by 40-ms depolarizing voltage steps in 5 mV increment (inset voltage protocol). Note that only the first 15 ms of the current traces are shown. The biophysical properties of Q1494A, Q1494E, Q1494L, and Q1494 channels relative to WT are shown in B and Table 1.

**Table 1.** Biophysical characteristics of the engineered and pathogenic Nav1.2 variants relative to wild-type (WT).

| Variant | I<sub>Na</sub> density at<br>–10 mV<br>(pA/pF) | V<sub>0.5,act</sub><br>(mV) | V<sub>0.5,inact</sub><br>(mV) | I<sub>Na-P</sub><br>at –10 mV<br>(% of peak) | τ recovery<br>(ms) |
| --- | --- | --- | --- | --- | --- |
| **WT** | 458.3 ± 55 | –17.56 ± 0.42 | –51.70 ± 0.96 | 1.06 ± 0.29 | 1.19 ± 0.03 |
| **n** | 22 | 22 | 22 | 22 | 18 |
| **P value** | – | – | – | – | – |
| **N1662D** | 95.2 ± 17<sup>****</sup> | –17.24 ± 0.44<sup>NS</sup> | –39.56 ± 1.3<sup>****</sup> | ND | 0.54 ± 0.02<sup>****</sup> |
| **n** | 16 | 16 | 16 |  | 14 |
| **P value** | <0.0001 | 0.999 | <0.0001 |  | <0.0001 |
| **Q1494A** | 212.7 ± 28<sup>**</sup> | –13.39 ± 0.47<sup>****</sup> | –36.75 ± 2.1<sup>****</sup> | ND | 0.51 ± 0.04<sup>****</sup> |
| **n** | 6 | 6 | 6 |  | 5 |
| **P value** | 0.048 | <0.0001 | <0.0001 |  | <0.0001 |
| **Q1494E** | 195.8 ± 54<sup>**</sup> | –16.01 ± 0.72<sup>NS</sup> | –36.15 ± 1.3<sup>****</sup> | ND | 0.61 ± 0.04<sup>****</sup> |
| **n** | 6 | 6 | 6 |  | 6 |
| **P value** | 0.002 | 0.463 | <0.0001 |  | <0.0001 |
| **Q1494L** | 103.0 ± 14<sup>****</sup> | –15.28 ± 0.74<sup>*</sup> | –43.02 ± 1.4<sup>****</sup> | ND | 0.69 ± 0.03<sup>****</sup> |
| **n** | 7 | 7 | 7 |  | 7 |
| **P value** | <0.0001 | 0.044 | <0.0001 |  | <0.0001 |
| **Q1494K** | 38.3 ± 7.9<sup>****</sup> | –9.03 ± 0.58<sup>****</sup> | –43.22 ± 0.46<sup>****</sup> | ND | 1.01 ± 0.08<sup>**</sup> |
| **n** | 9 | 9 | 8 |  | 8 |
| **P value** | <0.0001 | <0.0001 | <0.0001 |  | 0.0038 |
| **F1651C** | 461.5 ± 60<sup>NS</sup> | –16.12 ± 0.59<sup>NS</sup> | –38.92 ± 0.53<sup>****</sup> | 4.86 ± 0.57<sup>****</sup> | 0.67 ± 0.02<sup>****</sup> |
| **n** | 7 | 7 | 7 | 7 | 6 |
| **P value** | >0.999 | 0.482 | <0.0001 | <0.0001 | <0.0001 |
| **M1501V** | 265.4 ± 51<sup>*</sup> | –10.81 ± 0.61<sup>****</sup> | –42.63 ± 0.68<sup>****</sup> | 4.04 ± 0.79<sup>**</sup> | 0.84 ± 0.04<sup>****</sup> |
| **n** | 7 | 7 | 6 | 7 | 7 |
| **P value** | 0.032 | <0.0001 | 0.0001 | 0.0028 | <0.0001 |
| **M1501T** | 275.2 ± 27<sup>NS</sup> | –18.18 ± 0.58<sup>NS</sup> | –46.37 ± 0.54<sup>NS</sup> | 3.63 ± 0.96<sup>*</sup> | 0.69 ± 0.01<sup>****</sup> |
| **n** | 6 | 6 | 6 | 6 | 6 |
| **P value** | 0.076 | 0.996 | 0.007 | 0.021 | <0.0001 |
| **L1657P** | 109.8 ± 24<sup>****</sup> | –16.51 ± 0.84<sup>NS</sup> | –44.82 ± 3.7<sup>**</sup> | ND | 0.62 ± 0.03<sup>****</sup> |
| **n** | 7 | 7 | 7 |  | 6 |
| **P value** | <0.0001 | 0.844 | 0.0036 |  | <0.0001 |
| **P1658S** | 140.6 ± 22<sup>***</sup> | –14.34 ± 0.57<sup>***</sup> | –42.62 ± 0.68<sup>****</sup> | 5.91 ± 0.99<sup>****</sup> | 0.59 ± 0.02<sup>****</sup> |
| **n** | 9 | 9 | 9 | 9 | 7 |
| **P value** | <0.0001 | 0.0002 | <0.0001 | <0.0001 | <0.0001 |
| **A1659V** | 361.3 ± 50<sup>NS</sup> | –17.00 ± 0.63<sup>NS</sup> | –46.66 ± 0.60<sup>*</sup> | 5.64 ± 0.43<sup>****</sup> | 0.77 ± 0.05<sup>****</sup> |
| **n** | 10 | 10 | 10 | 10 | 9 |
| **P value** | 0.562 | 0.992 | 0.026 | <0.0001 | <0.0001 |

Data are represented as mean ± SEM. The statistically significant differences between WT and mutant Nav1.2 channels were determined using one-way ANOVA, followed by Dunnett's post-hoc test (\*P < 0.05, \*\*P < 0.01, \*\*\*P < 0.001, and \*\*\*\*P < 0.0001); ND, not determined; NS, statistically not significant difference compared to WT.
Abbreviations: I<sub>Na</sub>, sodium current; I<sub>Na-P</sub>, persistent sodium current; n, number of independent experiments; τ recovery, time constant of recovery from fast inactivation; V<sub>0.5,act</sub>, membrane potential for half-maximal activation; V<sub>0.5,inact</sub>, membrane potential for half-maximal inactivation.

### Inter-helical hydrogen bonds between N1662 and Q1494 are essential for fast inactivation

We hypothesised that the hydrogen bonds between the N1662 and Q1494 residues are critical determinants of the fast inactivation gating mechanism. To this end, we mutated the Q1494 residue to amino acids with neutral, acidic, or basic side chains and examined the mutant channels biophysically. We predicted that the substitution of the neutral, polar Q1494 residue with a neutral, nonpolar A or L residue would destabilize or abolish the fast inactivated state, consistent with the absence of hydrogen bonds between N1662 and A1494 or L1494. However, the substitution of Q1494 to acidic E or basic K residue should still permit the formation of a single hydrogen bond between N1662 and E1494 or K1494 and enable the development of fast inactivation to some extent.

The Q1494A and Q1494L mutations substantially prevented fast inactivation during depolarizing test pulses and resulted in biophysical properties that were very similar to those of N1662D variant. The Q1494E mutant exhibited slowly decaying currents, whereas the Q1494K retained significant inactivation (Figures 2B and 2C). Relative to wild-type, all four engineered mutations resulted in decreased current densities, faster recoveries of the inactivated fraction, and large depolarizing shifts of the inactivation curve (Figures 2B and 2C, Table 1), but had limited effects on activation, resulting in small shifts of the activation curve (Figures 2B and 2C, Table 1). These results demonstrate that hydrogen bond interactions between the N1662 and Q1494 residues represent critical molecular prerequisites for the development of fast inactivation in Na_v_1.2 channels.

### Molecular dynamics (MD) simulations reveal a reduced stability of the inactivated state in mutant channels

To confirm the structural basis underlying the disrupted fast inactivation in N1662D channels, we conducted MD simulations on the inactivated Na_v_1.2 structure. In over five replicates of 1 µs simulations, we assessed recovery from the fast-inactivated state by observing the relative stability of the DIII-IV linker and the binding of the IFM motif to its pocket.

The N1662D mutant showed pronounced differences in simulations compared to wild-type, displaying spontaneous dissociations of the DIII-IV linker within 1 µs in 3 out of 5 replicates, whereas the IFM motif of the wild-type channel remained bound across all replicates (Figures 3A-C, and SI Appendix Movie S1 and Movie S2). Distances between the centre of mass of IFM motif residues and the centre of mass of its binding site residues were significantly different between wild-type and the N1662D mutant (Figure 3D), suggesting that the mutation perturbs binding of the IFM motif. In wild-type, the ambifunctional hydrogen bonding configuration between Q1494 and N1662 was maintained throughout simulations (Figure 3E), contributing to stabilization of the bound DIII-IV linker in the inactivated state. Hydrogen bond donor interactions between N1662 and Q1494 were present in almost 100% in simulations of the wild-type. Notably, N1662 forms hydrogen bond donor interactions with the F1489 backbone (IFM motif residue), which likely further increases the binding stability of the DIII-IV linker. The mutation to D1662 results in loss of hydrogen bond donor interactions, due to lack of the nitrogen atom, and contributes to instability in binding of the linker (Figure 3E).

**Figure 3.**
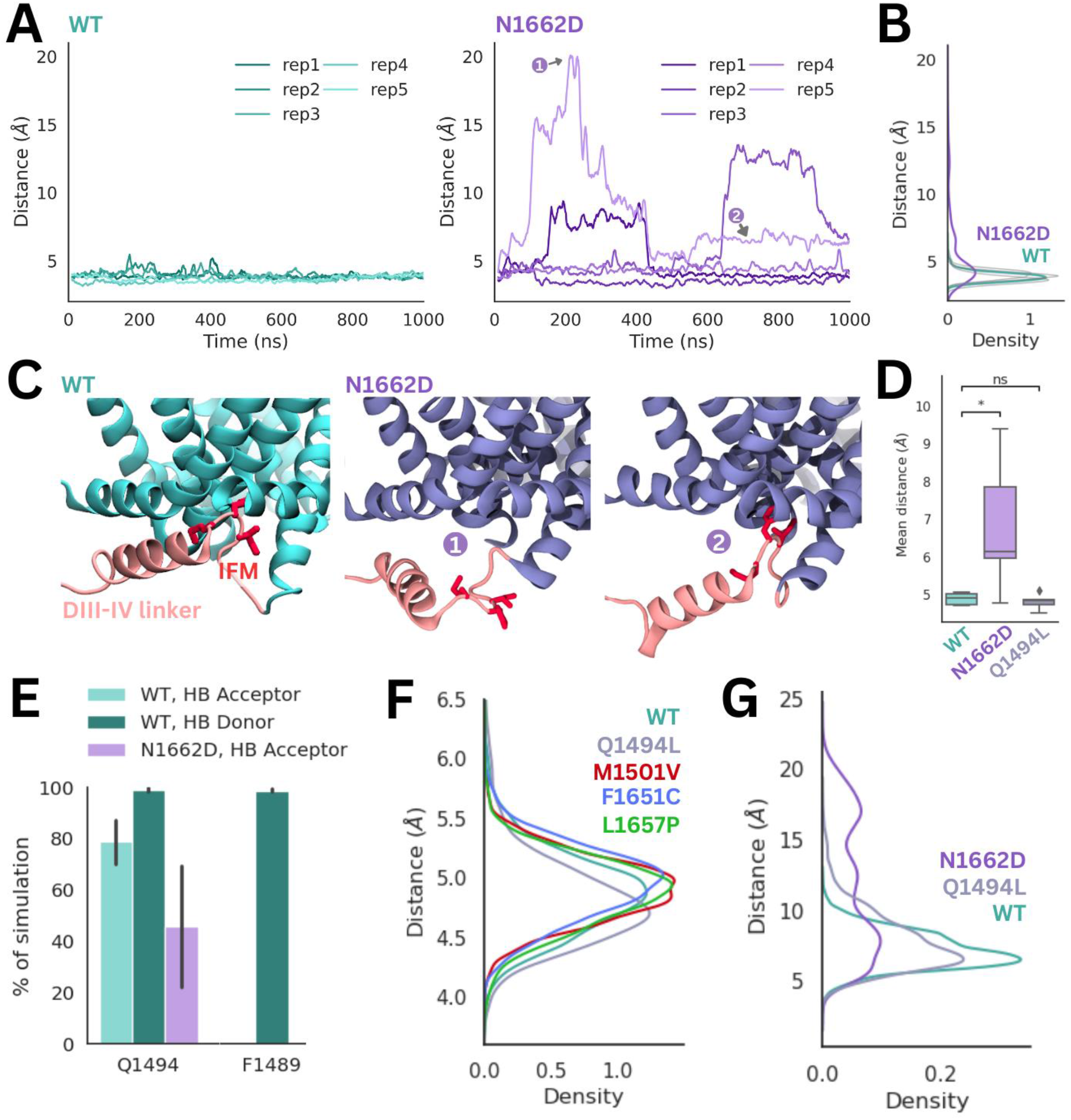
Stability of IFM-motif binding affected by different mutations compared to wild-type (WT) shown in molecular dynamics (MD) simulations. **(A)** Distance between the IFM motif and its binding pocket (defined as the center of mass of F1489 and the center of mass of residues 1659, 1663, and 1769) determined across the time course of each MD trajectory for WT (cyan) and N1662D (purple). Each of the five replicates plotted as a separate line. **(B)** Density plots showing differences in the distributions of distance between IFM motif and its binding pocket for WT and N1662D. **(C)** Representative snapshots taken from the trajectories showing the stably bound IFM motif in WT and the dissociation of the IFM motif and DIII-IV helix in N1662D at different timepoints (marked as 1 and 2). **(D)** The distance between the IFM motif and its receptor were averaged for each of the five replicates for WT, N1662D and Q1494L and shown as a boxplot. This distance significantly increased in the N1662D (P = 0.044) and was unchanged in Q1494L (P = 0.57). **(E)** Percentage of simulation where hydrogen donor and acceptor interactions were detected between residue 1662 and surrounding residues in WT (cyan) compared to N1662D (purple). **(F)** Distributions of distance between IFM motif and its binding pocket for other variants, Q1494L (grey), M1501V (red), F1651C (blue) and L1657P (green), were very similar to WT, as seen in unbiased MD simulations. **(G)** REST2 simulations were able to capture dissociation of the IFM in both N1662D and Q1494L.

To investigate contributions of Q1494 to stabilization of IFM motif binding, we simulated the Q1494L mutant. Surprisingly, unlike N1662D, Q1494L was very similar to wild-type with the IFM motif remaining stably bound across replicates (Figure 3D, 3F). Although the Q1494L mutation also disrupts the 1662-1494 hydrogen bonding network, the N1662 can still form a hydrogen bond with F1489, similar to WT (Figure 3E), thus reducing the propensity for IFM motif dissociation.

Given that the Q1494L mutation likely has a more subtle influence on the stability of the IFM motif than N1662D that was challenging to capture in within the timescale of our 1 µs unbiased MD simulations, we used an enhanced sampling method (REST2) to better examine stability of the IFM motif in the two mutants. For N1662D, the distribution of the distance between the IFM motif and its receptor shows two peaks greater than 10 Å, indicating a greatly reduced binding affinity, compared to wild-type (Figure 3G). The Q1494L variant shows a more subtle effect where the distances are slightly greater than the wild-type, suggesting a decreased binding stability of the IFM (Figure 3G). Overall, this highlights how different mutations in the DIII-IV linker can have varying effects on IFM binding and therefore inactivation, with N1662D having a stronger effect than Q1494L.

Apart from unbinding of the IFM motif, dissociation of other regions of the DIII-IV linker was also captured in the N1662D equilibrium simulation. Notably, the helical portion of the DIII-IV linker is observed to fluctuate in its orientation and move away from the pore gate (Figure 3C, right two panels). Thus, we hypothesized that residues in the DIII-IV linker, upstream of the IFM motif, are also responsible for binding to the S4-S5DIV linker and are also important for the inactivation mechanism. Overall, MD simulations demonstrate that interactions between N1662 and Q1494 play an important role in the spatial organization of the DIII-IV linker and binding of the IFM motif to its receptor site.

### N1662D I_Na_ amplitude dependence of the membrane voltage responses in axon initial segment (AIS) hybrid neurons

It is plausible that in the axon initial segments of patients carrying non-inactivating heterozygous Na_v_1.2 variants such as N1662D, firing activity is modulated by the expression level of the mutant Na_v_1.2 channel protein. To evaluate the impact of the N1662D channels on neuronal excitability, we characterized the voltage responses of the AIS hybrid cell model incorporating wild-type or N1662D I_Na_ in dynamic action potential clamp (DAPC) experiments (Figure 4A). Hybrid neurons incorporating wild-type I_Na_ showed firing activity with a characteristic bell-shaped input-output relationship (Figure 5J). To mimic the reduced expression level of N1662D channel relative to wild-type (Figure 2A), the inward peak amplitude of the implemented N1662D I_Na_ was systematically reduced. For example, a 5-fold lower N1662D expression relative to WT corresponded to a 0.2 external I_Na_ fraction (∼80 pA N1662D I_Na_). Hybrid neurons incorporating N1662D I_Na_ alone, did not fire action potentials during the step stimuli but showed sustained depolarization of the membrane potential due to the non-inactivating N1662D I_Na_ (Figures 4C and 4D, and SI Appendix, Figure S1). Sustained V_m_ depolarization could be elicited with a range of N1662D I_Na_ amplitudes and its magnitude was positively correlated with the implemented N1662D I_Na_ fraction (Figure 4B).

**Figure 4.**
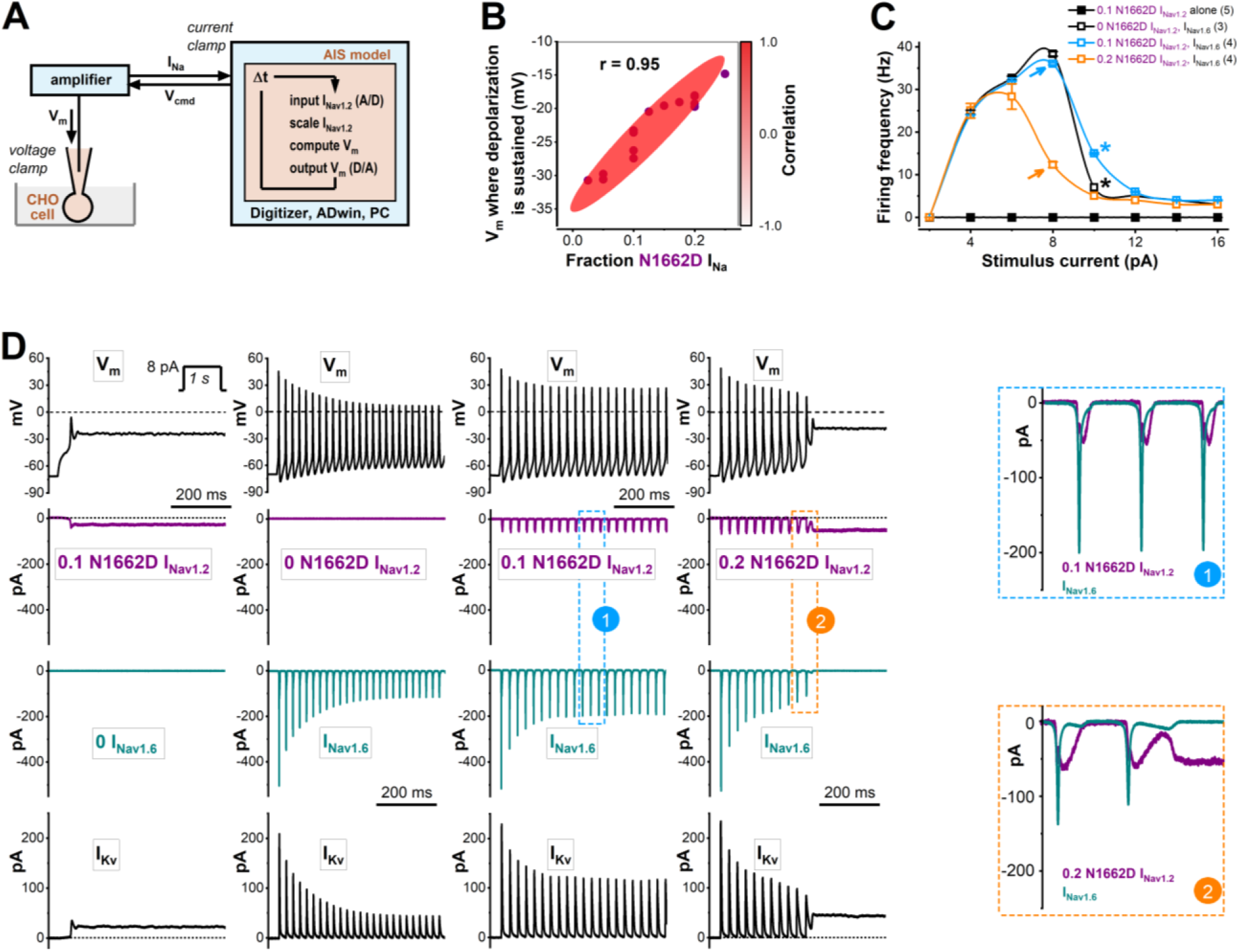
Excitability of hybrid neurons in the presence of wild-type (WT) or N1662D Na_v_1.2 currents. Dynamic action potential clamp (DAPC) experiments implementing. **(A)** Schematic representation of the dynamic action potential clamp (DAPC) configuration, consisting of a mammalian (CHO) cell transiently expressing WT or N1662D Na_v_1.2 channels and an axon initial segment (AIS) model. The membrane potential (V_m_) is calculated in real-time (using a time step, Δt, of 140 kH for updating the input and output values) and applied as a voltage clamp command to the CHO cell membrane to elicit WT or N1662D Na_v_1.2 current (I_Na_). This ‘external’ I_Na_ is scaled and implemented in the model neuron. Unless mentioned otherwise, the in silico Na_v_1.6 conductance is set to zero. **(B)** Relationships between the magnitude of the implemented N1662D (I_Na_ fraction) and the mean V_m_ where sustained depolarization develops. The shaded red area represents a strong positive correlation (r = 0.95); the number of individual experiments, n = 14. **(C)** Input-output relationships in the hybrid model neuron incorporating external N1662D I_Na_ alone (I_Nav1.2_, filled black squares), in silico wild-type Na_v_1.6 current alone (I_Nav1.6_, open black squares), I_Nav1.6_ and 0.1 external N1662D I_Nav1.2_ (open blue squares), or I_Nav1.6_ and 0.2 external N1662D I_Nav1.2_ (open orange squares). Relative to hybrid neurons incorporating 0.2 N1662D I_Nav1.2_ and I_Nav1.6_, neurons with an increased N1662D I_Nav1.2_ fraction show higher firing activity (orange and blue arrows, respectively). Note that firing activity can be higher in hybrid cells implementing 0.1 N1662D I_Nav1.2_ and I_Nav1.6_ relative to I_Nav1.6_ alone (blue and black stars, respectively). **(D)** Relationships between the V_m_ and selected membrane current components of the hybrid model neuron, showing representative V_m_ changes (top traces) and associated scaled external input N1662D I_Nav1.2_, in silico I_Nav1.6_ (downward deflections), and in silico voltage-gated potassium currents (I_Kv_, upward deflections). From left: N1662D I_Nav1.2_ alone (purple traces) elicits sustained depolarization. Note the associated sustained non-inactivating I_Nav1.2_ and the depressed I_Kv_ (black traces). In silico I_Nav1.6_ alone (dark cyan traces) is associated with sustained firing. In the presence of 0.1 or 0.2 I_Nav1.2_, in silico I_Nav1.6_ differentially enhances or reduces firing. Dashed lines indicate the zero-mV membrane potential (V_m_) level; dotted lines indicate the zero-pA current level; inset step current stimulus; the first 500 ms of the 1 s traces of V_m_ and current traces are shown.. See relationships between the inward currents associated with action potential firing from the boxed areas marked with ‘1’ and ‘2’.

**Figure 5.**
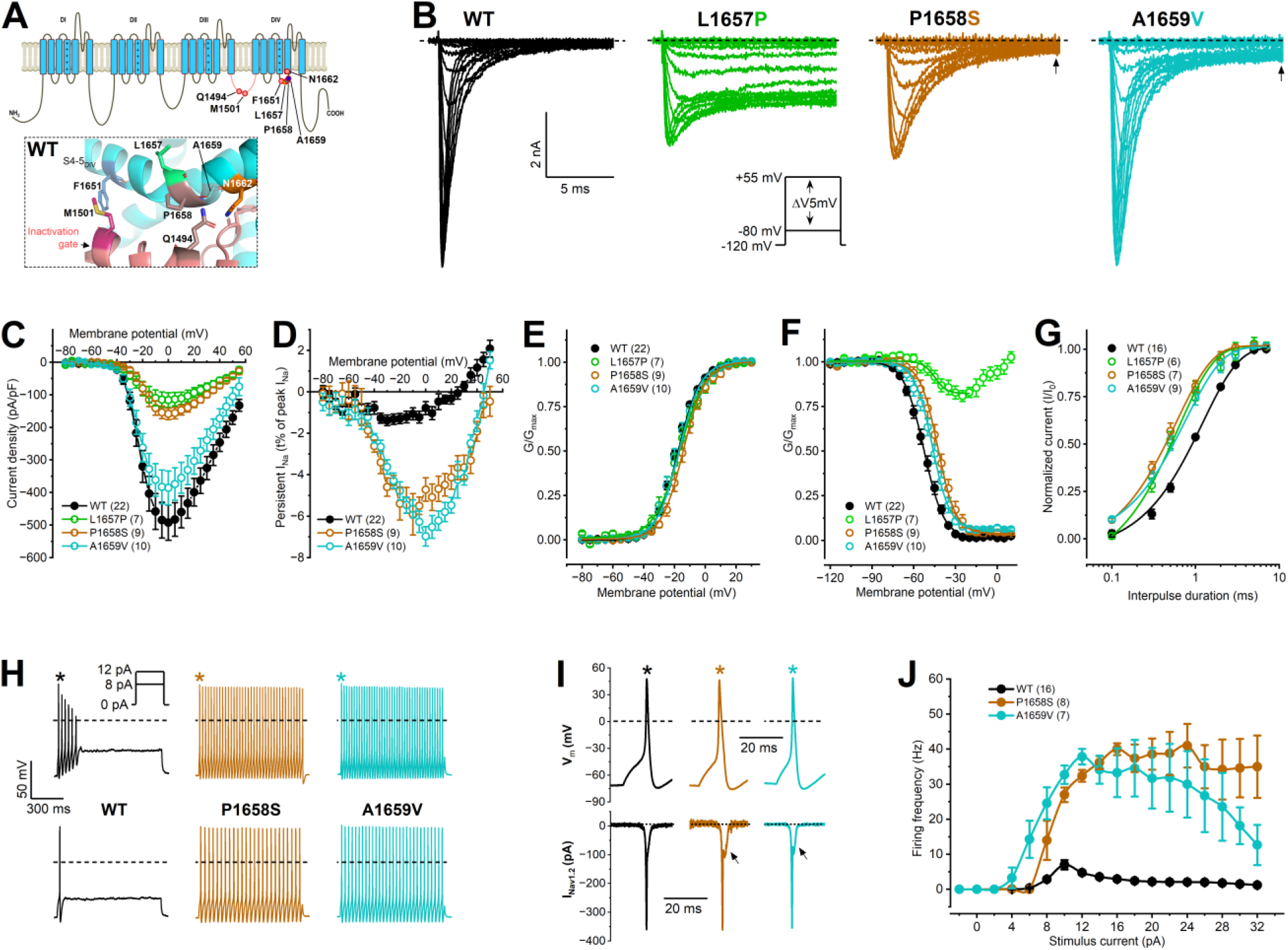
Biophysical properties and functional impact of the wild-type (WT) and pathogenic L1657P, P1658S, and A1659V variants. **(A)** Schematic representation of the assessed mutations relative to the N1662 residue. Zoomed-in cartoon representation of the S4-5D_IV_ linker structure in the WT. For additional details see Figure 1A. **(B)** Representative WT and mutant I_Na_ traces elicited from a holding potential of −120 mV by 40-ms depolarizing (conditioning) voltage steps in 5 mV increment (inset voltage protocol). The L1657P mutation produces non-inactivating current, whereas the P1658S and A1659V mutations result in a large persistent current relative to WT (arrows). Dashed lines indicate zero current level; only the first 14 ms of the current traces elicited in the voltage range between −80 and +10 mV are shown. **(C)** Current density-voltage relationships. **(D)** Persistent inward I_Na_-voltage relationships. **(E)** Voltage dependence of activation **(F)** Voltage dependence of steady-state inactivation. **(G)** Recovery from fast inactivation. Data in E-G were obtained and fitted as described in Figure 1. See parameters of the fits and statistical evaluation in Table 1. **(H)** Representative action potential traces in hybrid neurons incorporating transiently expressed WT, P1658S, or A1659V Na_v_1.2 currents in DAPC experiments. Traces elicited using 8 or 12 pA step stimuli are shown; dashed lines indicate zero V_m_ level. **(I)** Action potentials (corresponding to the action potentials marked by asterisks in H) and associated WT (black), P1658S (brown), or A1659V (turquoise) I_Nav1.2_ traces on expanded time scale; arrows indicate the increased persistent current component relative to WT;; dotted lines indicate zero current level. **(J)** Input-output relationships. Firing frequencies of mutants relative to WT were assessed using two-way ANOVA followed by Dunnett’s post-hoc test (*P < 0.05). Data are mean ± SEM; n, number of individual experiments in parentheses.

Hybrid neurons incorporating in silico I_Nav1.6_ alone elicited repetitive firing showing accommodation (Figures 4C and 4D). We varied the amplitude of the external N1662D Na_v_1.2 current (I_Nav1.2_) in a hybrid neuron in the absence or presence of ‘co-expressed’ in silico Na_v_1.6 current (I_Nav1.6_). Surprisingly, implementing 0.1 N1662D I_Nav1.2_ and in silico I_Nav1.6_ caused increased excitability, less accommodation, action potentials with increased amplitude, and more efficient action potential repolarization relative to I_Nav1.6_ alone (Figures 4C and 4D). However, a further increase of the non-inactivating I_Nav1.2_ fraction to 0.2 in the presence of I_Nav1.6_ led to a decrease in firing and transitioning of the membrane potential to a sustained depolarized value (Figures 4C and 4D). These experiments also demonstrate that membrane potential changes are associated with dynamic and complex changes of the current components in the hybrid neuron, contributing to the shifts in balance between inward (I_Na_) and outward potassium (I_Kv_) currents (Figure 4D).

### Na_v_1.2 missense mutations affecting the fast inactivation gating machinery predominantly result in hyperexcitability

The S4-5_DIV_ and DIII-IV linkers may undergo relatively large movements during fast inactivation gating, thus missense mutations affecting residues located in these regions are expected to severely impact channel function. We evaluated the biophysical characteristics and the impact on neuronal excitability of several DEE-associated missense mutations located in the inactivation gate (M1501T, M1501V) or S4-5DIV linker (F1651C, L1657P, P1658S, A1659V). The clinical features and treatment response of individual patients are summarized in SI Appendix.

The L1657P, P1658S, and A1659V mutations are localized near the N1662 residue (Figure 5A). Voltage clamp experiments revealed reduced current density and a depolarizing shift of the activation curve for P1658S, whereas the current densities and the activation curve for A1659V remained unchanged relative to wild-type (Figures 5C and 5E, Table 1). For both P1658S and A1659V, the inactivation curves shifted towards depolarized voltages, and the recoveries from fast inactivation were faster (Figure 5F and 5G, Table 1). Although P1658S and A1659V retained fast inactivation, these variants exhibited a very large inward persistent current (I_Na-P_) relative to wild-type, corresponding to ∼6 % of the transient peak I_Na_ (Figure 5D, Table 1). I_Na-P_ can have a considerable impact on excitatory neuronal function, influencing repetitive firing and facilitating hyperexcitability (30). Relative to wild-type, the P1658S or A1659V I_Na_ resulted in a significantly higher firing activity in DAPC experiments, associated with an increased I_Na-P_ during the repolarization phase (Figure 6H-J).

The L1657P variant resulted in a large, non-inactivating I_Na_, (Figure 5B). The biophysical properties of the L1657P variant were similar to those of N1662D (Figures 5C-G, Table 1). It is likely that the L to P mutation breaks or kinks the S4-5_DIV_ α-helix and disrupts the formation of the hydrogen bonds that are needed for maintaining the stability of the protein structure. In DAPC experiments, hybrid neurons incorporating L16757P I_Na_ alone showed sustained depolarization. In the presence of in silico I_Nav1.6_, a relatively small-amplitude external L16757P I_Na_ led to a transient increase of firing activity followed by sustained membrane depolarization, similar to N1662D I_Na_ (SI Appendix, Figure S2).

The 3D structure of the Na_v_1.2 channel suggests that in addition to hydrogen bond interactions shown in Figures 1 and 2, additional interactions may also contribute to the stability of the surface exposed, highly mobile inactivation gate. The residues M1501 and F1651, located on the DIII-IV linker and the S4-S5_DIV_ linker, respectively (Figure 5A, and SI Appendix, Figure S3), are thought to form a critical aromatic-sulfuric interaction, given that the F1651A mutation is known to result in a slow and incomplete fast inactivation (11). Mutations of the M1501 or F1651 residue may affect fast inactivation by destabilizing the binding of the IFM motif, however the effect may be less pronounced relative to those caused by N1662D. In MD simulations of the M1501V and F1651C mutations, we observed some instability between the DIII-IV linker and the S4-S5_DIV_ linker, however dissociation of the IFM motif could not be revealed with the IFM motif consistently bound throughout 1 µs simulations (Figure 3F). SI Appendix Figure S3 shows the biophysical consequences of the pathogenic F1651C, M1501V, and M1501T mutations and their impact on neuronal firing. As expected, these mutations also severely affected the voltage dependence and the recovery from fast of inactivation and increased the inward I_Na-P_ component relative to wild-type. DAPC experiments provided a rapid prediction of the neuron-scale phenotypic consequences of these three variants, resulting in increased action potential firing of the hybrid neuron relative to wild type (SI Appendix, Figure S3). The gain-of-function effects are consistent with the reported severe clinical phenotypes of these variants.

## DISCUSSION

In sodium channels, the binding of the hydrophobic IFM motif to the inactivation receptor site represents an essential conformation change leading to fast inactivation (1–3, 31). Inactivation can be prevented by destroying the inactivation gate using internally perfused pronase (32), preventing the outward movement of the S4_DIV_ using extracellular site 3 toxins (33), or mutating residues of the IFM motif or the inactivation receptor (10, 11). In this study, we demonstrate that fast inactivation of the Na_v_1.2 channel can also be prevented by missense mutations associated with early-onset DEE.

Recent high-resolution structure data demonstrate that the IFM motif binds to a receptor site located on pore regions located outside of the pore-lining helices (15, 17). The rate of inactivation is largely aligned with the relatively slow movement of S4_DIV_, after which the IFM motif may bind through a weakly voltage-dependent step to its receptor (4). Despite the proposed molecular models of fast inactivation gating (12, 15, 16), the details of the interactions between the residues involved are not fully understood.

### The influence of disrupted inactivation on channel function and neuronal excitability

Genetic diagnoses and functional-clinical evaluation of rare missense Na_v_1.2 variants revealed that pathogenic mutations can occur at various channel regions and can lead to gain-of-function or loss-of-function (34). Electrophysiological evaluation of Na_v_1.2 variants confirmed gain-of-function produces early-onset DEE (26, 35). Here, we performed voltage clamp and dynamic action potential clamp analyses to evaluate the biophysical and neurophysiological consequences of missense Na_v_1.2 variants associated with early-onset DEE. Seven mutations in channel sequences significantly involved in fast inactivation gating were studied.

Five variants demonstrated the expected gain-of-function. The F1651C, P1658S, A1659V, M1501V, and M1501T variants shifted the voltage dependence of steady-state inactivation to more positive membrane potentials and resulted in fast recovery from inactivation. The mutant currents displayed enhanced inward I_Na-P_ with an amplitude between 3 and 6 % of the transient inward peak I_Na_ because of incomplete or delayed fast inactivation. In DAPC experiments, each variant caused hyperexcitability when implemented in an excitatory hybrid neuron model.

Remarkably, the N1662D and L1657P variants did not demonstrate gain-of-function, but instead resulted in non-inactivating currents, highlighting a novel mechanism for *SCN2A*-related early-onset DEE. Given that sodium current inactivation represents a crucial aspect of neuronal excitability regulation, the lack of inactivation can lead to a prolonged depolarized state and affect the ability of neurons to fire action potentials effectively. DAPC experiments demonstrate that in the absence of other types of sodium conductances, N1662D or L1657P I_Na_ invariably results in the sustained depolarization of the hybrid neuron model.

However, it is important to note that both the N1662D and the L1657P variant showed reduced current densities relative to wild-type in transfected mammalian cells. This suggests that the impact on neuronal excitability may vary depending on several factors, including the variability in *SCN2A* allelic expression, the contribution of other neuronal sodium channels, and/or other cellular mechanisms. When implementing N1662D or L1657P I_Na_ with ∼10-fold reduced density relative to the enabled in silico Na_v_1.6 conductance, the mutant current triggers increased excitability in DAPC experiments. It is likely that this effect is due to the mutant current causing a relatively small depolarization of the membrane and/or contributing inward current during the activation of other types of neuronal sodium channels.

### Structural determinants of fast inactivation in Na_v_1.2 variants

Molecular dynamics (MD) simulations studies were performed using a Na_v_1.2 structure with an activated S4_DIV_ and IFM motif bound to its pore-module receptor (13) from which the IFM motif spontaneously dissociates in the N1662D mutant, reflecting a mutation-disrupted inactivation state. In agreement with the functional data, MD simulations revealed an increased distance between D1662 and Q1694 relative to wild type channel, indicating that the N1662D mutation reduces the affinity of the IFM motif to the DIVS4-S5 linker. The data also suggest that multiple hydrogen bond formation between the N1662 and Q1494 side chains is essential for the stability of the local DIII-IV linker region in the inactivated state. We hypothesise that instability of the linker is responsible for the accelerated rates of recovery from fast inactivation in the N1662D variant (Figure 2B) and contributes to overall disruption of the inactivation mechanism. This outcome substantiated mutagenesis experiments demonstrating that fast inactivation is also prevented in the Q1494A or Q1494L Na_v_1.2 channel variants, whereas the Q1494E or Q1494K variants enable an incomplete inactivation characterised by an enhanced I_Na-P_ relative to wild-type.

Although MD simulations were able to reveal significant differences in the distance between the IFM and its receptor pocket in the N1662D variant relative to wild-type, the differences for Q1494L were more subtle, only clearly showing in enhanced sampling MD simulations. Similarly, the respective distance for the L1657P variant was unchanged relative to wild-type (Figure 3F). It is likely that both Q1494L and L1657P allow some hydrogen bonds to remain and thus have a weaker influence on the affinity of the IFM motif to its receptor. The mutational effects of M1501V/T and F1651C are less clear. We do not see strong deficiency of fast inactivation in electrophysiology experiments, which is recapitulated in MD simulations where the IFM-motif does not dissociate. The M1501-F1651 interactions may have more complex structural implications that control the voltage dependence of inactivation.

### Challenges for treatment

The anti-seizure sodium channel blockers phenytoin, carbamazepine, and lamotrigine bind to receptor sites located inside the central cavity of the pore (36) and have a higher affinity to the fast inactivated state than to the resting state (37). In addition to these actions, the slow inactivation of the channel may be affected by phenytoin (38) and the voltage sensors may also be implicated in phenytoin binding (39). The lack of fast inactivation in N1662D and L1657P channels suggest that the affinity to common anti-seizure sodium channel blockers may be reduced. Indeed, the individual with the N1662D variant had little benefit from most sodium channel blockers used. However, given that neither the N1662D nor the L1657P mutation alters the voltage dependence of activation, it is also plausible that the sodium channel blocking effect of phenytoin is unaffected or relatively less affected. Supporting this latter hypothesis, in the patients with the L1657P variant (and to a lesser extent the individual with the N1662D variant), phenytoin treatment reduced seizures, although frequent seizures still persisted (SI Appendix). Future experiments are needed to clarify the effect of the commonly used anticonvulsants on non-inactivating sodium channel variants. Targeting the non-inactivating current with ranolazine (40) or using open channel blockers such as propafenone (16) may also be efficient.

In conclusion, we demonstrate that the N1662-Q1494 interaction is essential for maintaining the binding affinity of the IFM motif to its receptor in fast inactivated Na_v_1.2 channels. Our data have implications for elucidating fast inactivation of Na_v_1.2 channels and interpreting the impact of Na_v_1.2 variants on neuronal excitability. Non-inactivating Na_v_1.2 channel variants may result in a sustained depolarization, however the overall effect on neuronal excitability may depend on the expression level of the channel protein.

## MATERIALS AND METHODS

### SCN2A variants and patient data

Pathogenic *SCN2A* variants were identified through the *SCN2A* International Natural History Study (NHS) database (c.4984A>G, p.N1662D); Clinvar database (https://www.ncbi.nlm.nih.gov/clinvar/) (c.4502T>C, p.M1501T; c.4972C>T, p.P1658S); Simons Searchlight database (SSDb) (https://www.sfari.org/resource/simons-searchlight/) (c.4501A>G, p.M1501V); or referred through a network of collaborating clinicians (c.4970T>C, p.L1657P). The c.4952T>G, p.F1651C and c.4976C>T, p.A1659V variants have been reported previously (34, 41). The study was approved by the Human Research Ethics Committees of the Royal Children’s Hospital and Austin Health Melbourne, State Medical Association of Berlin, and the Simons Foundation. Written informed consent was obtained for the individuals with the L1657P and N1662D variants, whose clinical data is presented here.

### Na_v_1.2 channel variants

For all variants, the adult *SCN2A* isoform (42) was used as template. The N1662D, Q1494A, Q1494E, Q1494L, and Q1494K variants were synthesized using QuikChange mutagenesis kit (Agilent Technologies, Santa Clara, CA) with custom made forward (F) and reverse (R) primers (Bioneer Pacific) as follows: N1662D, 5’-ATGTCCCTTCCTGCGTTGTTTGACATCGGCCTC-3’ (F) and 5’-GAGGCCGATGTCAAACAACGCAGGAAGGGACAT-3’ (R); Q1494A, 5’-GGTCAAGACATTTTTATGACAGAAGAAGCGAAGAAATACTACAATGCAATGAAAAA-3’ (F) and 5’-TTTTTCATTGCATTGTAGTATTTCTTCGCTTCTTCTGTCATAAAAATGTCTTGACC-3’ (R); Q1494E, 5’-GTCAAGACATTTTTATGACAGAAGAAGAGAAGAAATACTACAATGCAATG-3’ (F) and 5’-CATTGCATTGTAGTATTTCTTCTCTTCTTCTGTCATAAAAATGTCTTGAC-3’ (R); Q1494L, 5’-CAAGACATTTTTATGACAGAAGAACTGAAGAAATACTACAATGCAATGAAA-3’ (F) and 5’-TTTCATTGCATTGTAGTATTTCTTCAGTTCTTCTGTCATAAAAATGTCTTG-3’; Q1494K, 5’-GTCAAGACATTTTTATGACAGAAGAAAAGAAGAAATACTACAATGCAATG-3’ (F) and 5’-CATTGCATTGTAGTATTTCTTCTTTTCTTCTGTCATAAAAATGTCTTGAC-3’ (R).

All sequences were verified and confirmed at the Australian Genome Research Facility (Melbourne, Victoria, Australia). The F1651C, M1501V, M1501T, L1657P, P1658S, and A1659V variants were custom made (Genscript, Piscataway, NJ, USA).

### Cell culture and Transfection

Chinese hamster ovary (CHO) cells were cultured and transiently co-transfected with wild-type or mutant Nav1.2 channel variant (5 μg) and enhanced green fluorescent protein (1 μg) using Lipofectamine 3000 kit (Thermo Fisher Scientific), as previously described (26). Cells were incubated at 37°C in 5% CO_2_ for 24 hours. Three to four days post-transfection, the cells were dissociated using TrypLE Express (Thermo Fisher Scientific Australia Pty Ltd) and plated on glass coverslips for electrophysiological recordings.

### Electrophysiology data and analysis

Sodium currents through Nav1.2 channel variants (I_Na_) were recorded using in in the whole-cell configuration of the patch-clamp technique using an Axopatch 200B amplifier (Molecular Devices, Sunnyvale, CA) as previously described (26, 43). The CHO cells were superfused with extracellular solution containing 145 mM NaCl, 5 mM CsCl, 2 mM CaCl_2_, 1 mM MgCl_2_, 5 mM glucose, 5 mM sucrose, and 10 mM Hepes (pH = 7.4 with NaOH), at a rate of ∼0.2 ml/min. Patch pipettes of ∼1.5 MΩ resistance were pulled from borosilicate glass capillaries (GC150TF-7.5, Harvard Apparatus Ltd.) and filled with intracellular (pipette) solution containing 5 mM CsCl, 120 mM CsF, 10 mM NaCl, 11 mM EGTA, 1 mM CaCl_2_, 1 mM MgCl_2_, 2 mM Na_2_ATP, and 10 mM Hepes (pH = 7.3 with CsOH). Current and potentials were low-pass filtered (cut-off frequency 10 kHz) and digitized at 50 kHz. Series resistance was compensated by ≥85%, and potentials were corrected for the estimated liquid junction potential. The leak and capacitive currents were corrected using a −P/4 pulse protocol, except when using steady-state inactivation and recovery from fast inactivation protocols. The voltage protocols assessing I_Na_ activation, steady-state inactivation, recovery from fast inactivation, persistent I_Na_ (I_Na-P_), and I_Na_ kinetics were described previously (43) and are shown as insets in the figures.

Dynamic action potential clamp (DAPC) recordings were performed by implementing scaled wild-type or A1329D I_Na_ in a biophysically realistic axon initial segment (AIS) model, as previously described (43). Unless specified otherwise, the virtual sodium conductance of the AIS model was set to zero, whereas the virtual K_v_ channel and the virtual Na_v_1.6 channel conductance values were set to gK_v_ = 2 (twice the original gK_v_) and gNa_v_1.6 = 0.4, respectively (43). Action potential firing was elicited using step current injections of 1 s duration, in 2-pA increments between −2 and +24 pA. In all DAPC experiments, the membrane potential (V_m_), stimulus current, in silico I_Kv_, in silico I_Nav1.6_, and I_Nav1.2_ were simultaneously recorded.

Current densities were determined by dividing the inward peak I_Na_ current amplitudes by cell capacitance (C_m_). The voltage-dependence of the activation was determined from a holding membrane potential (HP) of −120 mV, using depolarizing voltage steps of 40 ms duration in 5 mV increments in the voltage range between −80 and +55 mV. Current density values were plotted against membrane voltage to obtain current density–voltage relationships. Conductance (G) was determined according to the equation G = I/(V−V_rev_), where V_rev_ is the reversal potential for Na^+^.

Normalized conductance values (G/Gmax) were plotted against membrane potential to obtain activation curves. The voltage dependence of steady-state fast inactivation was determined from a HP of −120 mV using preconditioning voltage steps of 100 ms duration in 5 mV increments in the voltage range between −80 and +20 mV, followed by a 20-ms test pulse to −10 mV to test the availability of the sodium current. Activation and inactivation curves were fit using the Boltzmann equation as follows:

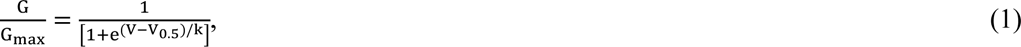

where V is the membrane voltage, V_0.5_ is the voltage for half-maximal activation or inactivation (V_0.5,act_ or V_0.5,inact_, respectively), and k is the slope factor. I_Na-P_ was determined after −P/4 leak correction, 40 ms after the onset of a depolarising voltage step and was expressed as percent of peak I_Na_ (43). I_Na-P_ was not determined for the engineered Na_v_1.2 variants (Q1494A, Q1494E, Q1494L, and Q1494K) and for the two pathogenic Na_v_1.2 variants (N1662D and L1657P). Q1494A, Q1494L, N1662D and L1657P exhibited non-inactivating inward current amplitudes ≥ 50% of the inward peak 40 ms after the onset of a depolarising voltage step.

The time constant of the recovery from fast inactivation was determined by measuring current amplitudes during paired-pulse (P1 and P2) depolarizations (43) and fitting a single exponential equation to data, as follows:

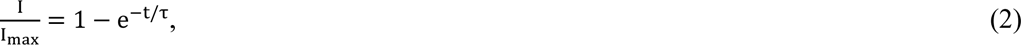

Where I_max_ is the current amplitude elicited by P1, I is the current amplitude elicited by P2, and t is the time between P1 and P2. For variants exhibiting a relatively large fraction of non-inactivating current (Q1494A, Q1494E, Q1494L, N1662D, and L1657P), the inactivated current fraction was first determined as the difference between the inward peak current amplitudes during P1 and P2 when using t = 0.1 ms. The recovery of the inactivated current fraction was plotted against t and a single exponential equation was fitted to the data as described above.

The firing frequency of the variants relative to wild-type was determined using Clampfit 10.7 software (Molecular Devices, Sunnyvale, CA). The mean firing frequency was plotted against the depolarizing stimulus current amplitude resulting in input-output relationships.

### 3D structural modelling

The voltage-gated Na_v_1.2 channel structure was downloaded from the Protein Data Bank (44) (PDB accession no. 6J8E) (13). The in silico mutations of Nav1.2 were introduced using Pymol 2.3.2 (Schroedinger LLC, New York, USA).

### Molecular dynamics (MD) simulations

Systems were set up for WT, N1662D, Q1494L, L1657P, M1501V and F1651C. The proteins were embedded within a pure 1-palmytoyl-2-oleoyl-sn-glycero-3-phosphatidylcholine (POPC) bilayer and solvated with 0.15 M NaCl solution using CHARMM-GUI (45). Mutations were introduced accordingly for each of the relevant systems. Systems were 140 Å along the plane of the membrane (x and y directions) and 120 Å in the z direction.

Simulations were conducted using Amber20 (46, 47), using the following forcefields: ffSB19 protein (48), Lipid22 (49), OPC water (50) and 12-6 ion parameters (51). All systems were equilibrated via minimizing, heating, pressurizing, a short 50 ns of protein backbone restrained simulation (at 5 kJ/mol), and gradual reduction of restraint over 24 ns. Each system was then simulated for five replicates of 1 µs each at 1 bar and 310 K, using the Monte Carlo barostat (52) and Langevin thermostat (53), respectively. Hydrogen mass repartitioning (54) was used to increase the timestep to 4 fs and hydrogen bonds were constrained via the SHAKE algorithm (55). Periodic boundary conditions and a 10 Å van der Waals cutoff were used. RMSD, RMSF and distance calculations were performed using MDAnalysis (56). Hydrogen bond interactions were assessed using ProLIF (57). Representative snapshots from MD trajectories were produced in Visual Molecular Dynamics (VMD) software (58).

Replica exchange with solute tempering (REST2) simulations were set up for WT, N1662D and Q1494L using the CHARMM-GUI Enhanced Sampler (59). Hamiltonians in the DIII-IV linker (residues 1486-1518) and sidechain atoms in the linker binding residues (1321-1330, 1476, 1479-1480, 1483, 1651-1663, 1769-1777) were scaled. 16 replicas were used at 310.15, 326.37, 343.20, 360.67, 378.75, 397.62, 417.08, 437.27, 458.17, 479.88, 502.42, 525.80, 550.06, 575.24, 601.42, 620.30 K respectively. Systems were equilibrated for 10 ns without scaled Hamiltonians, then each replica was equilibrated for 10 ns with scaled Hamiltonians. REST2 was run for 500 ns, with a timestep of 2 fs and attempting exchange every 1000 steps. Simulations were conducted with GROMACS 2022 (60) using the CHARMM36m (61) forcefield and TIP3P water (62), with the pressure set at 1 bar using the Parinello-Rahman barostat (63) and the global temperature set to 310 K using the Noose-Hoover thermostat (64). Analysis was conducted on the lowest temperature replica.

### Statistics

Data are presented as mean ± SEM. Electrophysiological data were analysed in Clampfit 9.2 (Molecular Devices). Statistical analyses were carried out using GraphPad Prism 9.0 (La Jolla, CA, USA) and Origin 2023 (Microcal Software Inc., Northampton, MA). One-way ANOVA followed by Dunnetts’ post hoc test was used to evaluate the properties of Na_v_1.2 channel variants relative to wild-type. Pearson’s correlation analysis was used to determine the strength of linear association between the fraction of N1662D I_Na_ and the V_m_ value showing sustained depolarization in DAPC experiments, and the correlation coefficient, r. Differences were considered statistically significant if P < 0.05; n, the number of individual experiments, are included in the figures and in Table 1.

Statistical analyses of the MD simulations were performed by first applying decorrelation analysis to extract uncorrelated data points using timeseries in pymbar python packages, then using the student’s t-test on the filtered data points via the scipy python package.

## AUTHOR CONTRIBUTIONS

GB, ET, BC and SP designed the study. GB designed mutagenesis primers, generated *SCN2A* constructs, carried out 3D structural modeling and electrophysiology studies. ET carried out molecular dynamics simulations. KBH, RKC and MW collected and interpreted clinical data. KK collated clinical data and designed mutagenesis primers for custom made *SCN2A* constructs. GB, ET, KBH, RKC, MW, BC, and SP wrote and edited the manuscript. All authors contributed to the editing of the manuscript.

## ACKNOWLEDGMENTS and FUNDING SOURCES

We thank the patients and their families for participating in our research. We acknowledge the Simons Foundation/Simons Searchlight database in providing data for this study. We appreciate obtaining access to phenotypic and genetic data on SFARI Base. Approved researchers can obtain the Simons Searchlight population dataset described in this study by applying at https://base.sfari.org. This study was supported by an Australian Research Council Centre of Excellence for Integrative Brain Function grant (CE14010007), National Health and Medical Research Council (NHMRC) program grant (10915693) to SP, Medical Research Future Fund Genomic Health Futures Mission Project Grant to GB, KBH, and SP, and project funding to the *SCN2A* Natural History study by RogCon, Inc. and Praxis Precision Medicines to KBH. SP is supported by an NHMRC Fellowship, and KBH by a clinician-scientist fellowship from the Murdoch Children’s Research Institute (MCRI). GB was partly funded by RogCon, Inc. ET is supported by a Research Training Program (RTP) Scholarship. The Florey Institute of Neuroscience and Mental Health and MCRI are supported by a Victorian State Government Operational Infrastructure Support Program. This work was supported by computational resources provided by the Australian Government through the National Computational Infrastructure (NCI) under the ANU and National Merit Allocation Schemes.

## Glossary

ASM: anti-seizure medication
CHO: Chinese hamster ovary
CTD: C-terminal domain
DAPC: dynamic action potential clamp
DEE: developmental and epileptic encephalopathy
LoF: loss-of-function
GoF: gain-of-function
MD: molecular dynamics
SCB: sodium channel block
RMSD: root mean square deviation
RMSF: root mean square fluctuation

## SUPPORTING INFORMATION

### Supporting Information Text

#### Patients

##### Non-inactivating variants

The **N1662D** variant, identified through the *SCN2A* International Natural History Study (NHS) database and reported in abstract form by Takacs et al, was identified in an individual born at 34 weeks gestation (24). This individual had early-infantile developmental and epileptic encephalopathy, with seizure onset on day one of life. Initial seizure types were focal and tonic, and EEG in the neonatal period demonstrated a burst-suppression pattern. Some initial reduction in seizure frequency was noted with phenobarbitone and phenytoin, although phenytoin was reported to increase sedation. Ultimately, seizures were resistant to all treatments tried, including sodium channel blocking (SCB) anti-seizure medication (SCB: lacosamide, oxcarbazepine, zonisamide, non-SCB: levetiracetam). She experienced frequent episodes of status epilepticus until initiation of the ketogenic diet (KD) at age 3 years of age. KD remains the most effective treatment and has significantly reduced ICU admissions. Her current epilepsy treatments are KD, lacosamide, oxcarbazepine, zonisamide and phenobarbitone. Her MRI Brain at ag 5 years showed severe cerebral and cerebellar atrophy, with diffusion restriction present in the bilateral symmetric medial temporal lobes, thalamus, and hippocampi. Her EEG at age six years showed a diffusely suppressed background without interictal epileptiform activity. This child continues to experience profound developmental delays, severe hypotonia and microcephaly, is gastrostomy-fed, and has a tracheostomy.

**L1657P**, has been identified in a male patient after a normal pregnancy and birth. Seizures started from day two of life onwards with frequent daily episodes of tonic seizures, spasms, and status epilepticus, with burst-suppression pattern on the EEG (consistent with EIDEE) and normal brain MRI. Treatment with phenobarbital, levetiracetam, vigabatrin or ketogenic diet showed no effect. With phenytoin treatment, seizures and EEG improved, but daily seizures persisted. Addition of valproate, bromide, or chloral hydrate had no effect on epilepsy. The child showed a severe global developmental disorder, an extreme severe form of spasticity, had no eye contact and little reaction to stimuli, and was fed with a PEG tube. The child deceased at age 20 months from pneumonia.

##### Variants showing gain-of-function

**P1658S**, identified through the Clinvar database (https://www.ncbi.nlm.nih.gov/clinvar/), has been described as a likely pathogenic variant.

**A1659V**, has been identified in a cohort of Vietnamese DEE patients and reported as pathogenic/likely pathogenic (41).

**M1501T** (Clinvar database), has been associated with early-onset DEE.

**M1501V**, identified through the Simons Searchlight database (SSDb) (https://www.sfari.org/resource/simons-searchlight/), has been reported in a male patient and described as a pathogenic de novo variant associated with early-infantile DEE.

**F1651C**, has been identified as a mosaic de novo mutation in a patient with early-onset DEE characterised by seizure onset at six weeks of age, with seizures classified as clonic and generalized tonic-clonic, showing multifocal spikes and episodes of status epilepticus. At three months, the patient became seizure free following phenytoin administration, however low levels of phenytoin resulted in relapse (34).

**Fig. S1.**
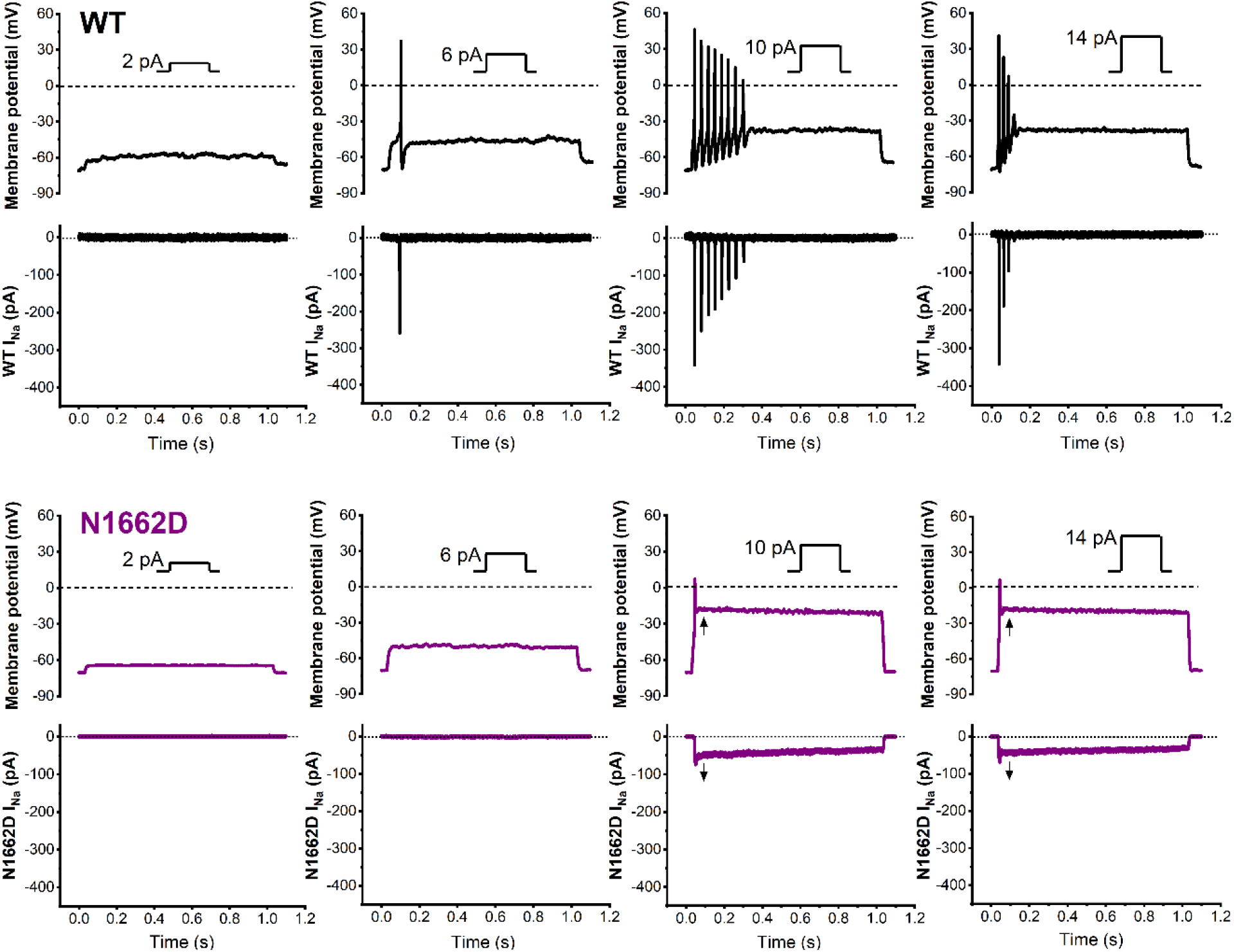
Different patterns of activity in the presence of heterologously expressed wild-type (WT) or N1662D Na_v_1.2 sodium current (I_Na_) in dynamic action potential clamp (DAPC) experiments. Hybrid neurons with WT I_Na_ show transient action potential firing activity in response to depolarizing step current stimuli of increasing amplitude (insets) (top, black traces, from left to right); see the input-output relationships in Figure 5J). The implemented external WT I_Na_ traces, associated with action potential firing, are shown as downward deflections (bottom, black traces). Relative to WT, hybrid neurons with N1662D I_Na_ show an initial passive and small-magnitude depolarization in response to depolarizing step current stimuli (top, purple traces, first two panels from left), followed by a switch to a sustained depolarized state (arrows), in response to higher amplitude depolarizing stimuli (10 and 14 pA) (top, purple traces, last two panels on the right). The associated external N1662D I_Na_ traces show no channel activity (I_Na_ = 0) in hyperpolarized state (bottom, purple traces, first two panels from left) or result in sustained inward current due to non-inactivating N1662D channels. Dashed lines indicate the zero-mV membrane potential level; dotted lines indicate the zero-pA current level.

**Fig. S2.**
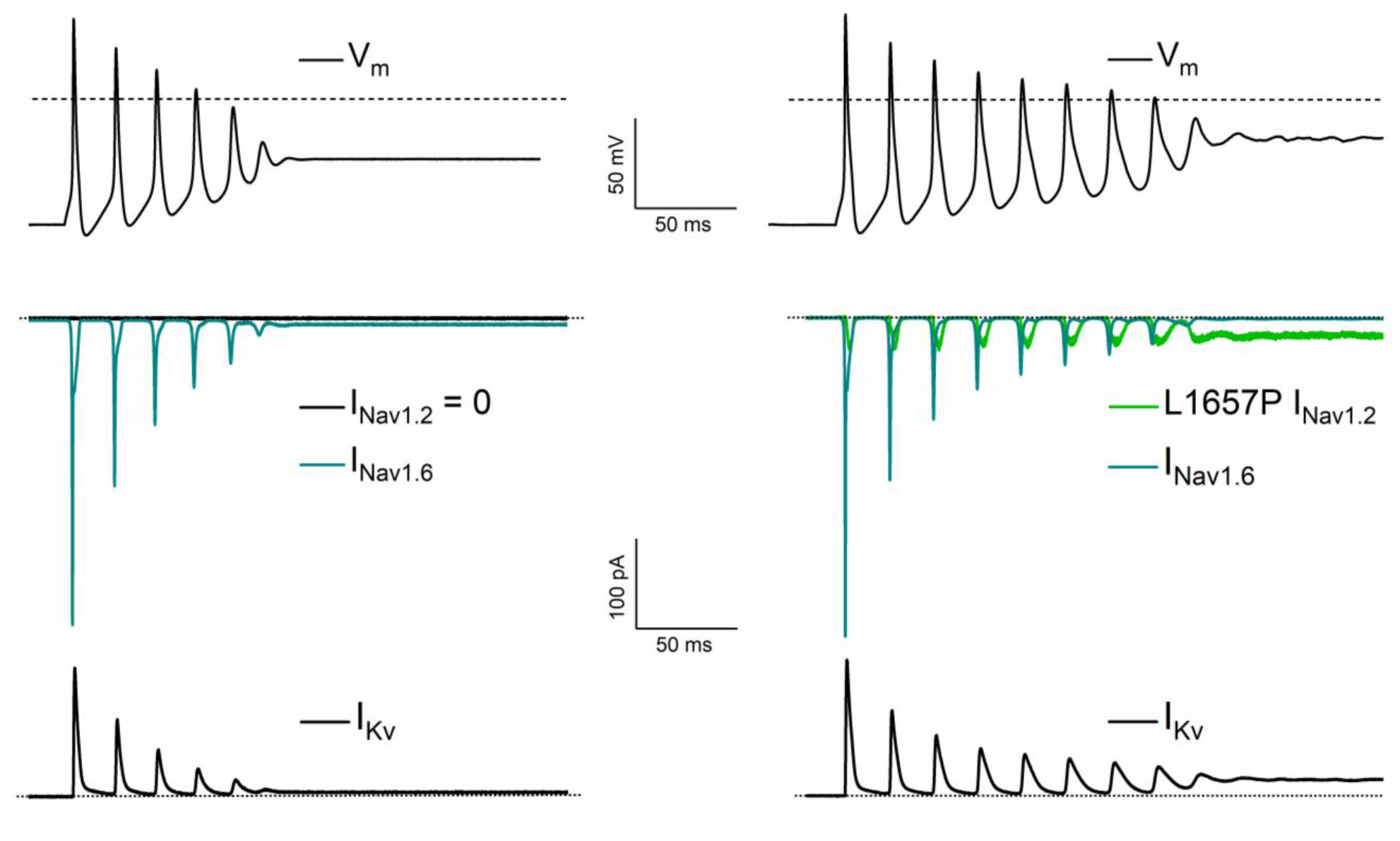
DAPC experiment demonstrating that a relatively small-amplitude external L16757P Na_v_1.2 current (I_Nav1.2_) increases action potential firing frequency in the presence of in silico Na_v_1.6 current (I_Nav1.6_). Action potential firing (top) in the absence (left) and presence (right) of L1657P I_Nav1.2_. The action potential associated external L1657P I_Nav1.2_ (green trace, middle right), in silico I_Nav1.6_ (dark cyan traces, middle, left and right) and in silico potassium currents (IK_v_, black traces, bottom, left and right) are shown. L1657P I_Nav1.2_ was scaled to a peak amplitude corresponding to approximately 10 % of that of inward peak I_Nav1.6_, resulting in a transient increase of the action potential firing transitioning into sustained depolarization of the membrane potential (V_m_). Note that only the first 300 ms of the 1 s traces of V_m_ and current traces, elicited upon the injection of a 12-pA stimulus current, are shown.

**Fig. S3.**
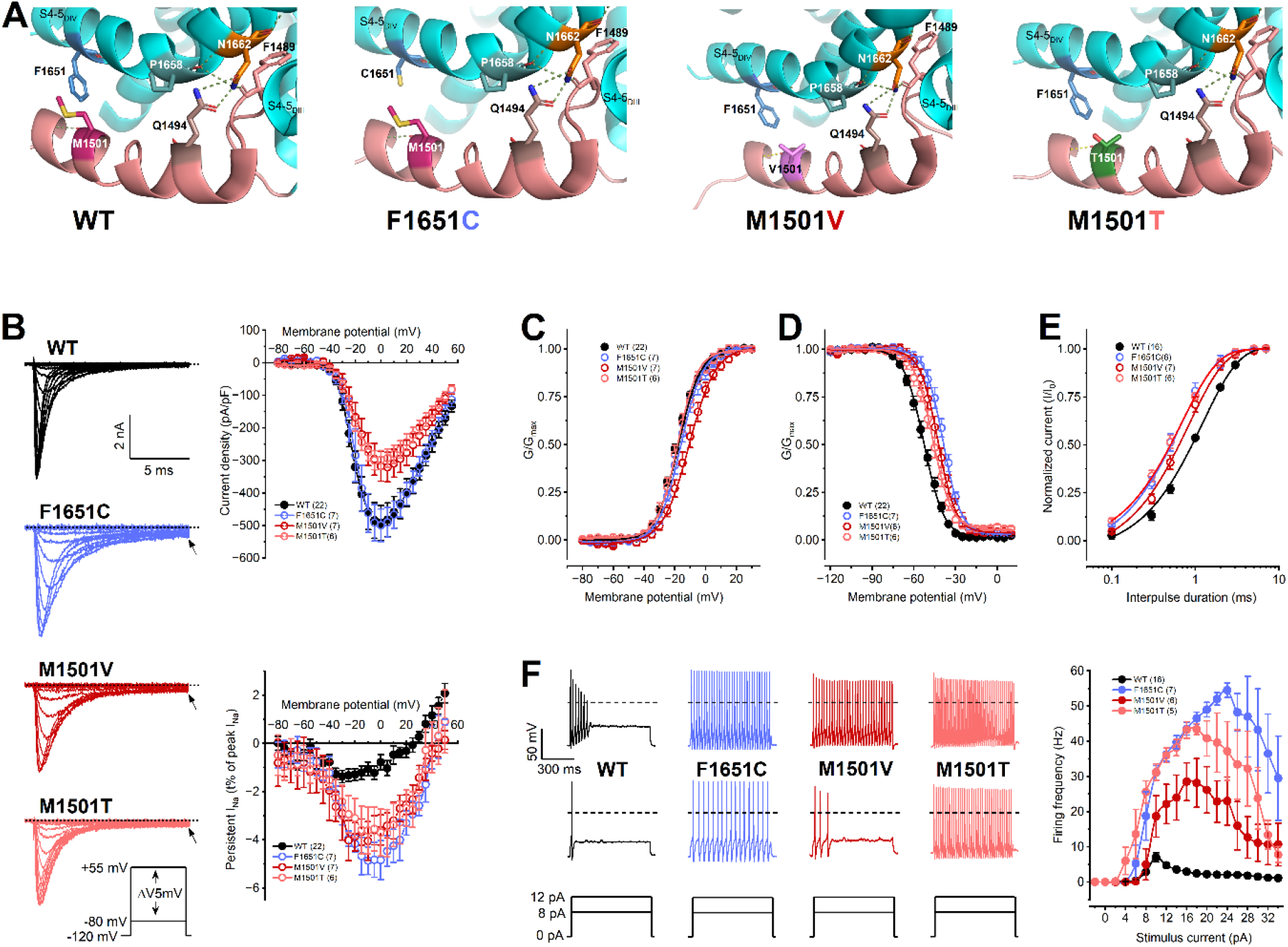
Biophysical properties and impact on hybrid neuron excitability of pathogenic F1651C, M1501V, and M1501T channels. (A) Zoomed-in views of the wild-type (WT) and mutant residues in segment 4-5 linker of domain IV (S4-5D_IV_) or the inactivation gate (a-helix in pink). Selected residues are represented as sticks, with the electronegative nitrogen amines (in blue), carbonyl oxygens (in red), hydroxyl group (red), and sulphur atom (yellow) of the amino acid side chains. Note the F1489 IFM motif residue in the inactivation gate. **(B)** Left: representative WT and mutant I_Na_ traces elicited in the voltage range between −80 and +10 mV. Top right: current density-voltage relationships. Top left: persistent inward sodium current (I_Na_)-voltage relationships (bottom); inset voltage protocol. Note the presence of persistent current with mutant channels (arrows) relative to WT. Dashed lines indicate zero current level. **(C)** Voltage dependence of activation **(D)** Voltage dependence of steady-state inactivation. **(E)** Recovery from fast inactivation. Data in C-E were fitted as described in Figure 1 (See Methods). See parameters of the fits and statistical evaluation in Table 1. **(F)** Representative action potential firing elicited by 8 and 12 pA step stimuli, and input-output relationships showing the effect of increasing stimulus strength on firing frequency in DAPC experiments. Dotted lines indicate zero membrane potential (V_m_) level. Firing frequencies relative to WT were assessed using two-way ANOVA followed by Dunnett’s post-hoc test (*P < 0.05). Data are mean ± SEM; n, number of individual experiments in parentheses.

**Movie S1 (separate file).** Stability of the DIII-IV linker and IFM motif binding in wild-type (WT) Na_v_1.2, captured in 1 µs of unbiased molecular dynamics simulation.

**Movie S2 (separate file).** Dissociation of the DIII-IV linker and IFM motif in the N1662D Na_v_1.2 variant, captured in 1 µs of unbiased molecular dynamics simulation.

